# Monoclonal Antibodies Capable of Binding SARS-CoV-2 Spike Protein Receptor Binding Motif Specifically Prevent GM-CSF Induction

**DOI:** 10.1101/2020.09.04.280081

**Authors:** Xiaoling Qiang, Shu Zhu, Jianhua Li, Ping Wang, Kevin J. Tracey, Haichao Wang

## Abstract

A severe acute respiratory syndrome (SARS)-like coronavirus (SARS-CoV-2) has recently caused a pandemic COVID-19 disease that infected more than 25.6 million and killed 852,000 people worldwide. Like the SARS-CoV, SARS-CoV-2 also employs a receptor-binding motif (RBM) of its envelope spike protein for binding the host angiotensin-converting enzyme 2 (ACE2) to gain viral entry. Currently, extensive efforts are being made to produce vaccines against a surface fragment of a SARS-CoV-2, such as the spike protein, in order to boost protective antibody responses. It was previously unknown how spike protein-targeting antibodies would affect innate inflammatory responses to SARS-CoV-2 infections. Here we generated a highly purified recombinant protein corresponding to the RBM of SARS-CoV-2, and used it to screen for cross-reactive monoclonal antibodies (mAbs). We found two RBM-binding mAbs that competitively inhibited its interaction with human ACE2, and specifically blocked the RBM-induced GM-CSF secretion in both human monocyte and murine macrophage cultures. Our findings have suggested a possible strategy to prevent SARS-CoV-2-elicited “cytokine storm”, and provided a potentially useful criteria for future assessment of innate immune-modulating properties of various SARS-CoV-2 vaccines.

**One Sentence Summary:** RBM-binding Antibodies Inhibit GM-CSF Induction.

## Introduction

Shortly after the 2003 outbreak of the severe acute respiratory syndrome (SARS) caused by a β-coronavirus (SARS-CoV) ^1^, the recent emergence and rapid spread of SARS-like coronavirus 2, SARS-CoV-2, has caused a pandemic COVID-19 that is catastrophically damaging human health. As of 1 September 2020, more than 25.6 million people have been infected, leading to more than 852,000 deaths in 216 countries (https://www.who.int/emergencies/diseases/novel-coronavirus-2019). Like the SARS-CoV ^1^, SARS-CoV-2 virus also employs its envelope spike (S) glycoproteins to recognize and bind a host cell surface receptor, the angiotensin-converting enzyme 2 (ACE2), to gain host cell membrane fusion and viral entry^2–8^. Structurally, the SARS-CoV-2 S protein contains a receptor-binding domain (RBD) that embraces a receptor-binding motif (RBM) in a “closed” configuration inaccessible by the host ACE2 receptor. Upon cleavage of the S protein by host proteases such as furin and the transmembrane protease/serine subfamily member 2 (TMPRSS2), the RBD undergoes a conformational change (from a “closed” to an “open” configuration) that enables the “exposure” of RBM to host cell receptors ^8–14^.

In the absence of effective therapies, vaccination has become a key option to boost adaptive antibody responses against SARS-CoV-2 infections. One approach is to use a surface fragment of a SARS-CoV-2, such as the spike (S) protein as antigens ^15^, in the hope that antibodies targeting the S protein may inhibit viral interaction with host ACE2 receptor to prevent viral entry ^15^. In patients infected by SARS-CoV or SARS-CoV-2, neutralizing antibodies targeting the RBD or RBM region of respective S proteins were found ^1, 3–7, 16–18^; and some of them indeed impaired RBD-ACE2 interaction ^17^ and viral entry ^4, 16^. Intriguingly, a previous study revealed that antibodies against different epitopes of SARS-CoV S protein exhibited divergent effects: antibodies targeting RBM (residue 471-503) conferred protection; whereas antibodies targeting epitopes (e.g., residue 597-603) outside of the RBM region worsen the outcomes ^19^. However, it was previously unknown how RBM-targeting antibodies would affect innate inflammatory responses to SARS-CoV-2 infections?

Recently, emerging evidence suggested that ACE2 might also be expressed on innate immune cells such as human peripheral mononuclear cells (PBMCs) ^20, 21^ and murine macrophage-like RAW 264.7 cells ^21^. Furthermore, human PBMCs produced several pro-inflammatory cytokines (e.g., TNF, IL-1β and IL-6) and chemokines (e.g., IL-8 and MIP-1β) in response to SARS-CoV S protein stimulation ^22^. However, it was previously unknown how RBM-binding monoclonal antibodies (mAbs) affect the SARS-CoV-2-elicited innate immune responses. In the present study, we sought to screen for mAbs capable of binding SARS-CoV-2 RBM, and determine how these RBM-binding mAbs affect the RBM-induced cytokine/chemokine production in monocyte and macrophage cultures.

## Results

### Generation of recombinant RBD and RBM protein fragments of SARS-CoV-2

To screen for monoclonal antibodies capable of binding the RBD or RBM region of SARS-CoV-2 spike protein (**Fig. 1A**), we generated recombinant RBD and RBM corresponding to residue 319-541 and residue 437-508 of SARS-CoV-2 spike (S) protein (**Fig. 1B**). These recombinant proteins were purified from insoluble inclusion bodies by differential centrifugation, urea solubilization, and histidine-tag affinity chromatography (**Fig. 1C**). Extensive washing of the immobilized recombinant RBD or RBM proteins with buffer containing 8.0 M urea effectively removed contaminating bacterial endotoxins. Subsequently, the purified RBD and RBM was dialyzed in a buffer supplemented with a reducing agent, Tris (2-carboxyethyl) phosphine (TCEP), to prevent excessive oxidation and cross-linking of the nine and two Cysteine (C) residues in RBD and RBM, respectively (**Fig. 1B**). As shown in **Fig. 1D**, amino acid sequence analysis revealed a high homology between a tyrosine (Y)-rich segment (YNYLYR) of SARS-CoV-2 RBM and the epitope sequence (NDAL<u>**YEYLR**</u>Q) of several monoclonal antibodies (mAbs) that we recently generated against human tetranectin (TN) ^23^, suggesting a possibility that some TN-binding mAbs might cross-react with SARS-CoV-2.

**Figure 1.**
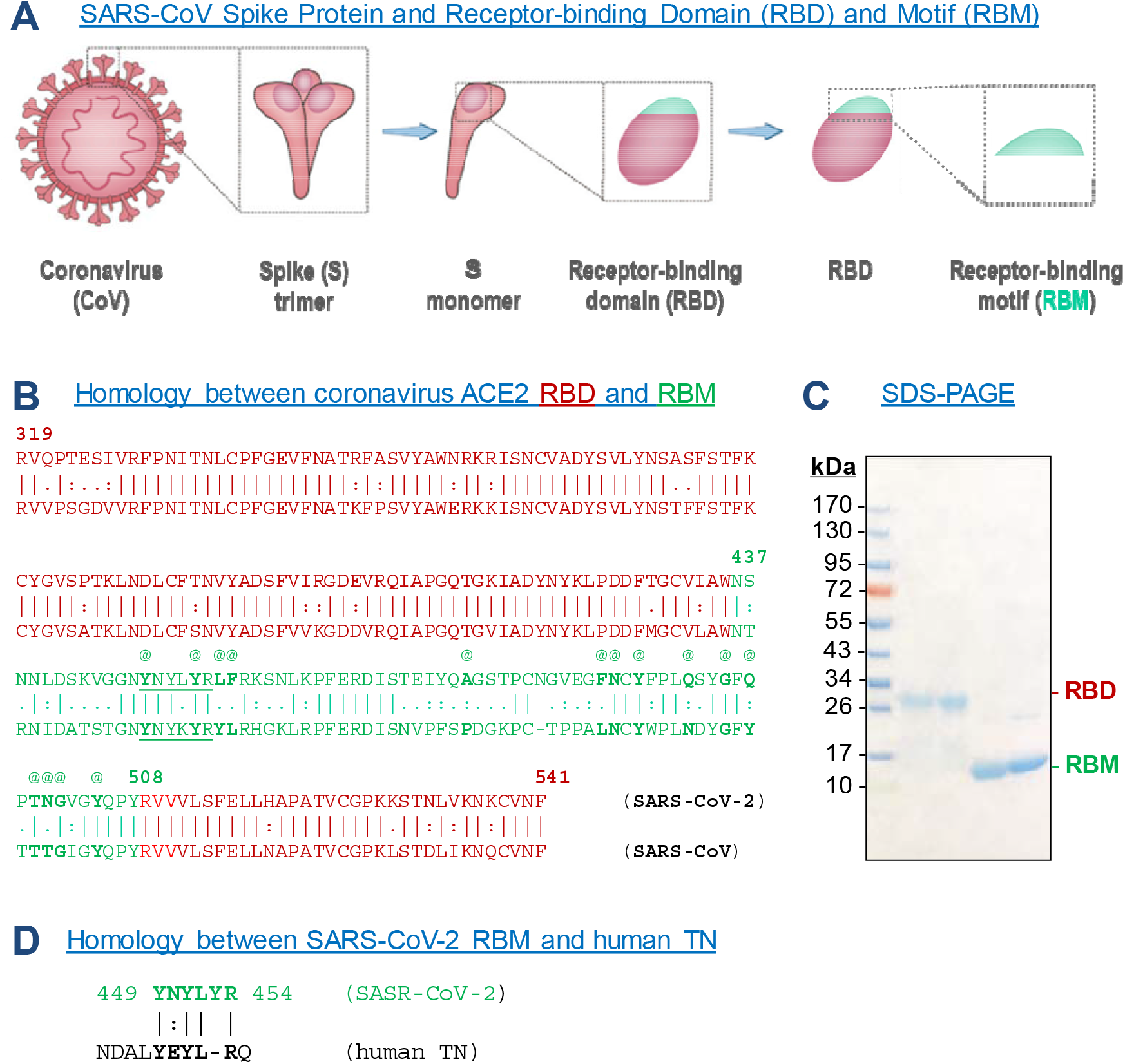
Generation of the ACE2 receptor-binding domain (RBD) and receptor-binding motif (RBM) of SARS-CoV-2 spike protein. **A**) Schematic diagram of SARS-CoV spike protein (S) and its ACE2 receptor binding domain (RBD) and motif (RBM). **B**) Amino acid sequence of RBD and RBM of SARS-CoV and SARS-CoV2. RBM sequence is denoted by text in green; “@”, denote residues in close contact with ACE2 (Lan, et al 2020). **C**) SARS-CoV-2 spike protein RBD and RBM corresponding to amino acids 319-541 and 437-508 with an N-terminal histidine tag were expressed in *E. coli* BL21 (DE3) pLysS cells, and purified by differential centrifugation of inclusion bodies, urea solubilization and histidine-tag affinity chromatography. **D**) SARS-CoV-2 RBM contains a sequence highly homologous to the epitope sequence (NDAL<u>YEYLR</u>Q) of several anti-TN monoclonal antibodies (mAbs).

### Recombinant RBM interacted with human ACE2 and some TN-binding mAbs

To evaluate the ACE2-binding properties of recombinant RBD or RBM, the extracellular domain of human ACE2 was immobilized to the NTA sensor chip, and recombinant RBD or RBM was applied as analytes at different concentrations to estimate the dissociation equilibrium constant (K_D_) using the Open SPR technique. Surprisingly, our recombinant RBD exhibited an extremely low affinity to the extracellular domain of human ACE2 (**Fig. 2A, upper panel**), with an estimated K_D_ of 161,000 nM. It was postulated that the cysteine-rich RBD was not likely re-folded into a “correct” conformation suitable for RBM-ACE2 interaction, because the high probability of “incorrect” disulfide cross-linking was factorially proportional to its high number of Cysteine residues. In contrast, the K_D_ for ACE2-RBM interaction ranged around 42.5 – 64.1 nM (**Fig 2A, lower panel; Fig. 2B**), regardless whether ACE2 or RBM was conjugated to the NTA sensor chip before respective application of RBM or ACE2 as analyte at different concentrations. Given the proximity between our estimated K_D_ for RBM-ACE2 interaction and the previously reported K_D_ (15 – 44.2 nM) for SARS-CoV-2 S protein-ACE2 interaction ^12, 18^, we concluded that the ACE2-binding property was well-preserved in our recombinant RBM. Therefore, we conjugated a highly purified recombinant RBM on an NTA sensor chip, and use it to screen for SARS-CoV-2 RBM-binding mAbs. In agreement with a homology between SARS-CoV-2 RBM and epitope sequence of several TN-specific mAbs (**Fig. 1D**), we found that 2 out of 3 mAbs capable of recognizing a homologous epitope sequence (NDALYEYLRQ) ^23^ exhibited a dose-dependent interaction with RBM (**Fig. 2C**), with an estimated K_D_ of 17.4 and 62.8 nM, respectively. This estimated K_D_ was comparable to that of other SARS-CoV-2 RBD-binding neutralizing antibodies (K_D_ = 14 −17 nM) recently isolated from COVID-19 patients ^24, 25^.

**Figure 2.**
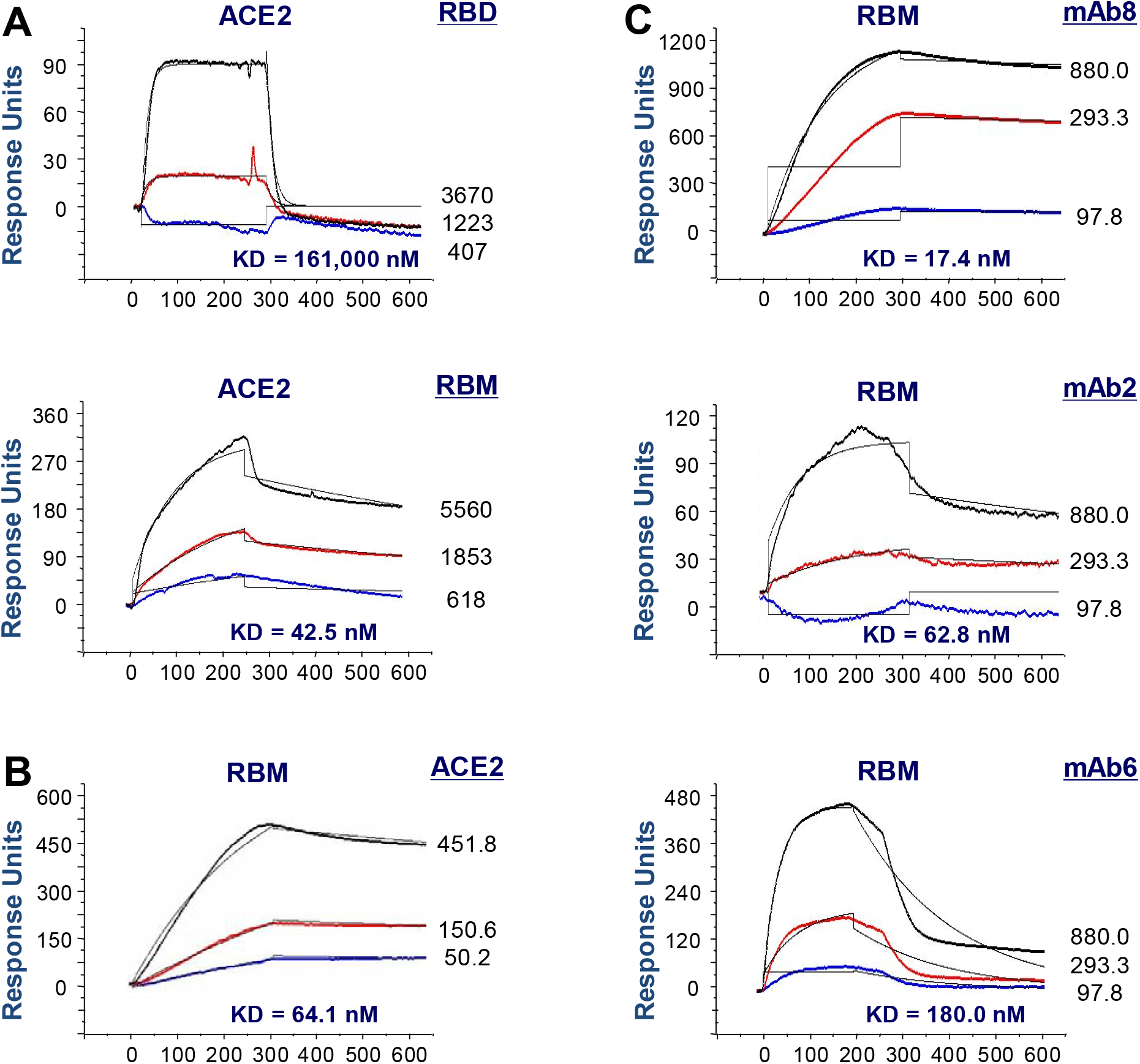
Recombinant SARS-CoV-2 RBM binds to human ACE2 receptor and some TN-reactive mAbs. **A**) Highly purified extracellular domain of human ACE2 was immobilized on a sensor chip, and recombinant RBD or RBM was applied as analyte at various concentrations to estimate the dissociation equilibrium constant (K_D_). **B, C**) Highly purified recombinant RBM was immobilized on the sensor chip, and recombinant human ACE2 (**Panel B**) or TN-specific mAbs (**Panel C**) were applied as analyte at various concentrations to assess the K_D_ for RBM-ACE2 (**Panel B**) or RBM-mAb (**Panel C**) interactions.

### RBM-binding mAbs competitively inhibited RBM-ACE2 interaction

We then tested whether pre-treatment of RBM-conjugated sensor chip with RBM-binding mAb competitively inhibited subsequent RBM-ACE2 interactions. When conjugated to a sensor chip, the recombinant RBM exhibited a dose-dependent interaction with the extracellular domain of human ACE2 (**Fig. 3A**), as well as a RBM-binding mAb (mAb8) (**Fig. 3B**). However, after pre-treatment with mAb8, the maximal response unit was markedly reduced from ~500 (**Fig. 3A**) to 175 (**Fig. 3C**) when ACE2 was applied as analyte to the RBM-coated sensor chip at identical concentrations. Meanwhile, the estimated K_D_ for RBM-ACE2 interaction was increased by an almost ten-fold from 13.3 nM (**Fig. 3A**) to 130.0 nM (**Fig. 3C**), suggesting that RBM-binding mAbs competitively RBM-ACE2 interactions.

**Figure 3.**
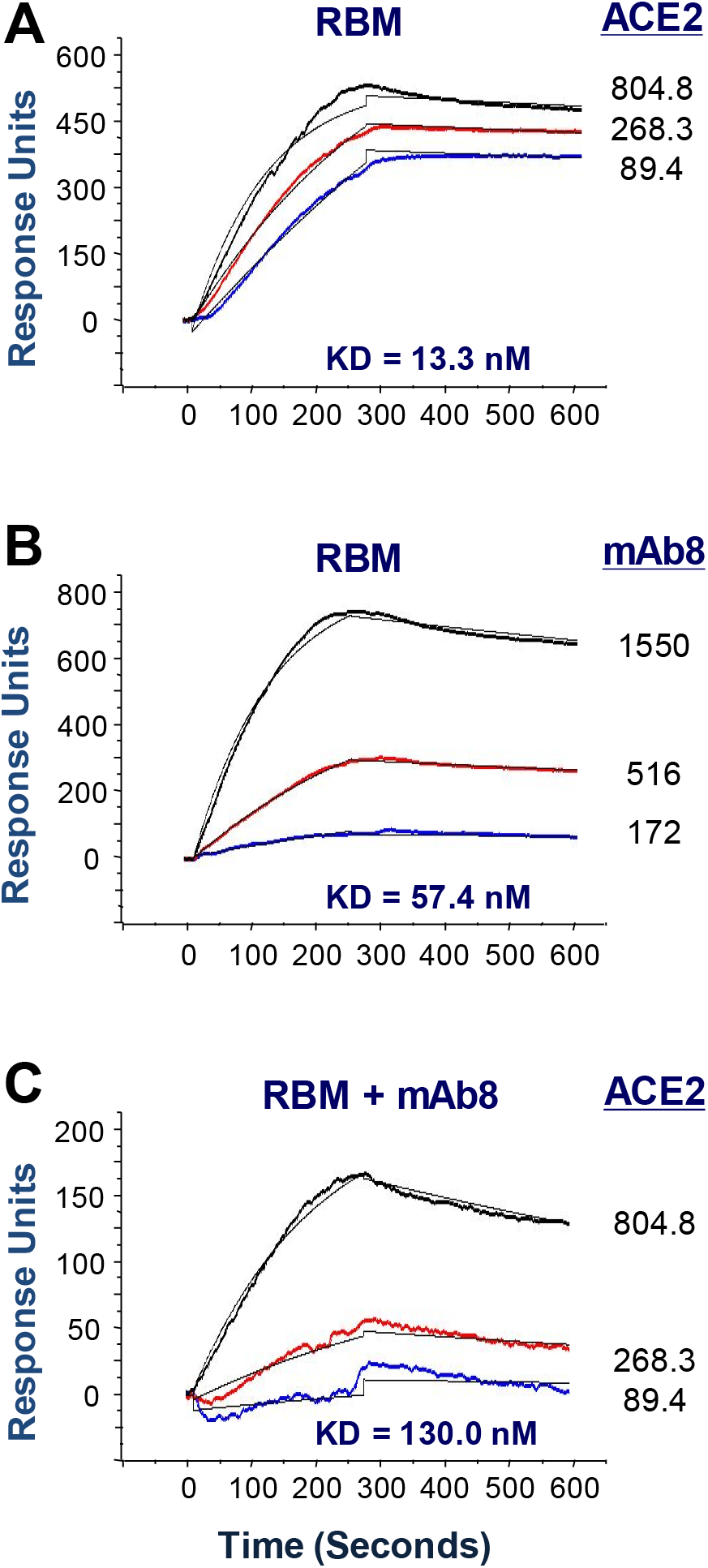
RBM-reacting mAb interferes with RBM-ACE2 interaction. **A)** Highly purified recombinant RBM was immobilized on the sensor chip, and recombinant human ACE2 was applied as analyte at various concentrations to assess the K_D_ for RBM-ACE2 interaction. **B**) After extensive washing, mAb8 was applied at indicated concentrations (**Panel B**), before ACE2 were re-applied to the RBM-conjugated sensor chip at identical concentrations (as in **Panel A**). The almost 10-fold increase in the KD (from 13.3 to 130.0 nM) and an almost 3-fold decrease (from 500 to 165) in the response units suggested that pre-treatment with mAb8 markedly inhibited RBM-ACE2 interaction.

### RBM-binding mAbs specifically blocked the RBM-induced GM-CSF secretion in primary human peripheral blood mononuclear cells (huPBMCs)

To examine the possible impact of RBM-binding mAbs on its immuno-stimulatory properties, human primary monocytes were stimulated with recombinant RBD or RBM in the absence or presence of RBM-binding mAbs (mAb8 and mAb2), and the levels of 42 different cytokines and chemokines were measured simultaneously by Antibody Arrays. In agreement with a previous report that SARS-CoV spike (S) protein stimulated human PBMCs to produce proinflammatory cytokines (e.g., IL-1β, IL-6, and TNF) ^22^, we observed a marked elevation of these three cytokines in the RBD- or RBM-stimulated human monocytes (**Fig. 4A, 4B**). In addition, both RBD and RBM also markedly stimulated the secretion of an anti-inflammatory cytokine (IL-10) and two chemokines (MIP-1δ and MCP-1) in parallel (**Fig. 4A, 4B**). Astonishingly, our highly purified RBM, but not RBD, also markedly induced the secretion of a myeloid growth factor, the granulocyte–macrophage colony-stimulating factor (GM-CSF) in human monocytes (**Fig. 4A, 4B**). However, the co-addition of two RBM-binding mAbs similarly and specifically blocked the RBM-induced secretion of GM-CSF (**Fig. 4A, 4B**) without affecting the RBM-induced release of other cytokines (e.g., IL-1β, IL-6, IL-10 and TNF) or chemokines (MIP-1δ and MCP-1).

**Figure 4.**
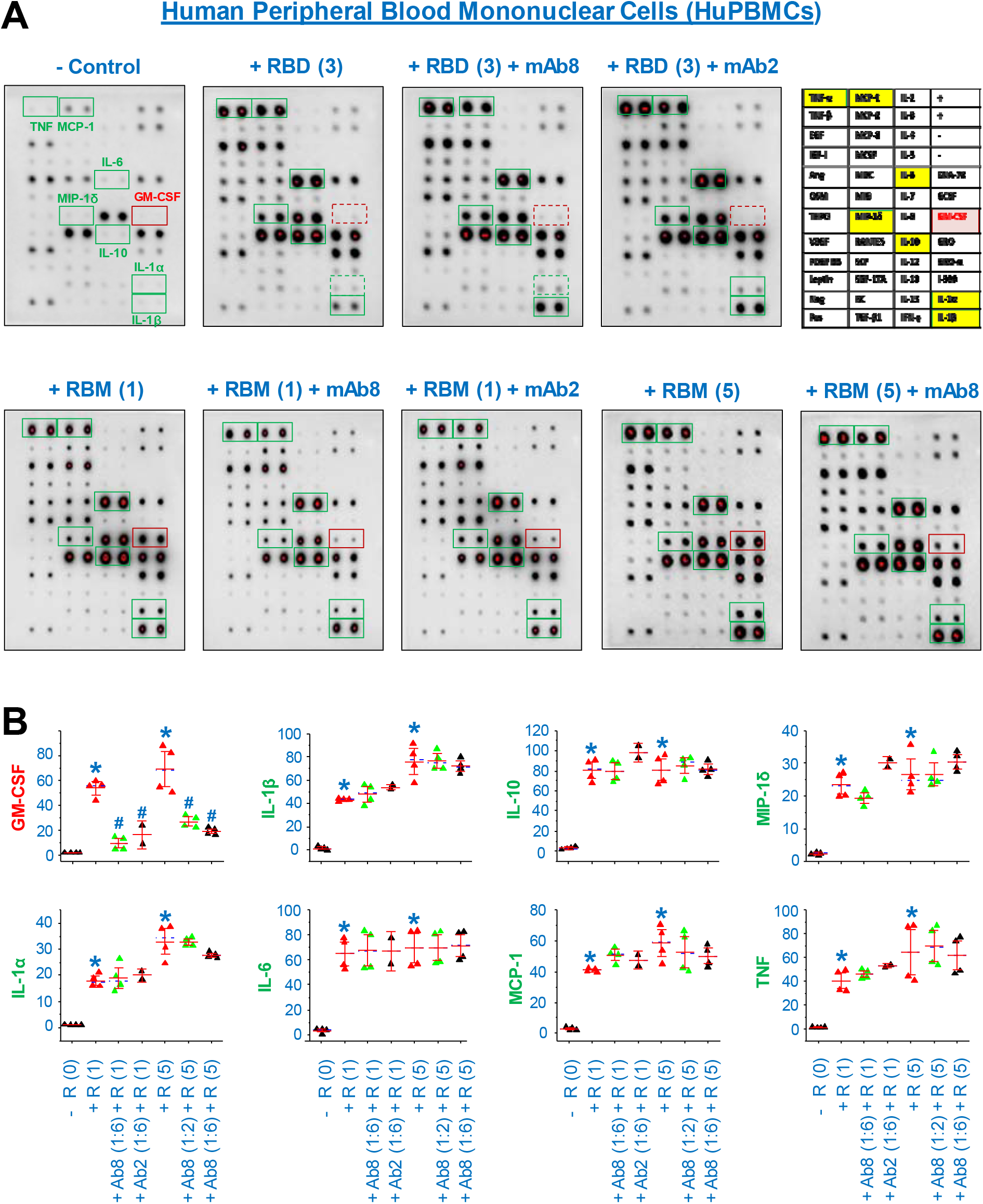
RBM-reactive mAbs abrogated the RBM-induced secretion of GM-CSF in human peripheral blood mononuclear cells (PBMCs). Human peripheral blood mononuclear cells (HuPBMCs) were isolated from blood of healthy donors, and stimulated with recombinant RBD (3.0 μg/ml) or RBM (1.0 or 5.0 μg/ml) in the absence or presence of RBM-binding mAbs (mAb8 or mAb2, at a molar ratio of 1:2 or 1:6). At 16 h post stimulation, the extracellular concentrations of 42 different cytokines and chemokines were determined by Cytokine Antibody Arrays (Panel A), and normalized by the positive controls (“+”) on respective membranes (Panel B). *, *P* < 0.05 versus negative controls (“-RBM” or “-R”); #, *P* < 0.05 versus positive controls (“+ RBM” or “+R”) at respective concentrations.

### RBM-binding mAbs also specifically blocked the RBM-induced GM-CSF secretion in murine macrophage-like RAW 264.7 cells

To further confirm the GM-CSF-inducing activities of SARS-CoV-2 RBM, we stimulated murine macrophage-like RAW 264.7 cells with highly purified RBM in the absence or presence of RBM-binding mAbs, and measured the extracellular levels of 62 different cytokines by Antibody Arrays. Compared with human monocytes, murine macrophages appeared to be less responsive to RBM stimulation, and released relatively fewer cytokines after stimulation (**Fig. 5A**). However, SARS-CoV-2 RBM still markedly elevated the secretion of TNF and GM-CSF in murine macrophage cultures (**Fig. 5A**). Similarly, two different RBM-binding mAbs selectively blocked the RBM-induced GM-CSF secretion in macrophage cultures (**Fig. 5A, 5B**) without affecting the RBM-induced TNF secretion (**Fig. 5A, 5B**).

**Figure 5.**
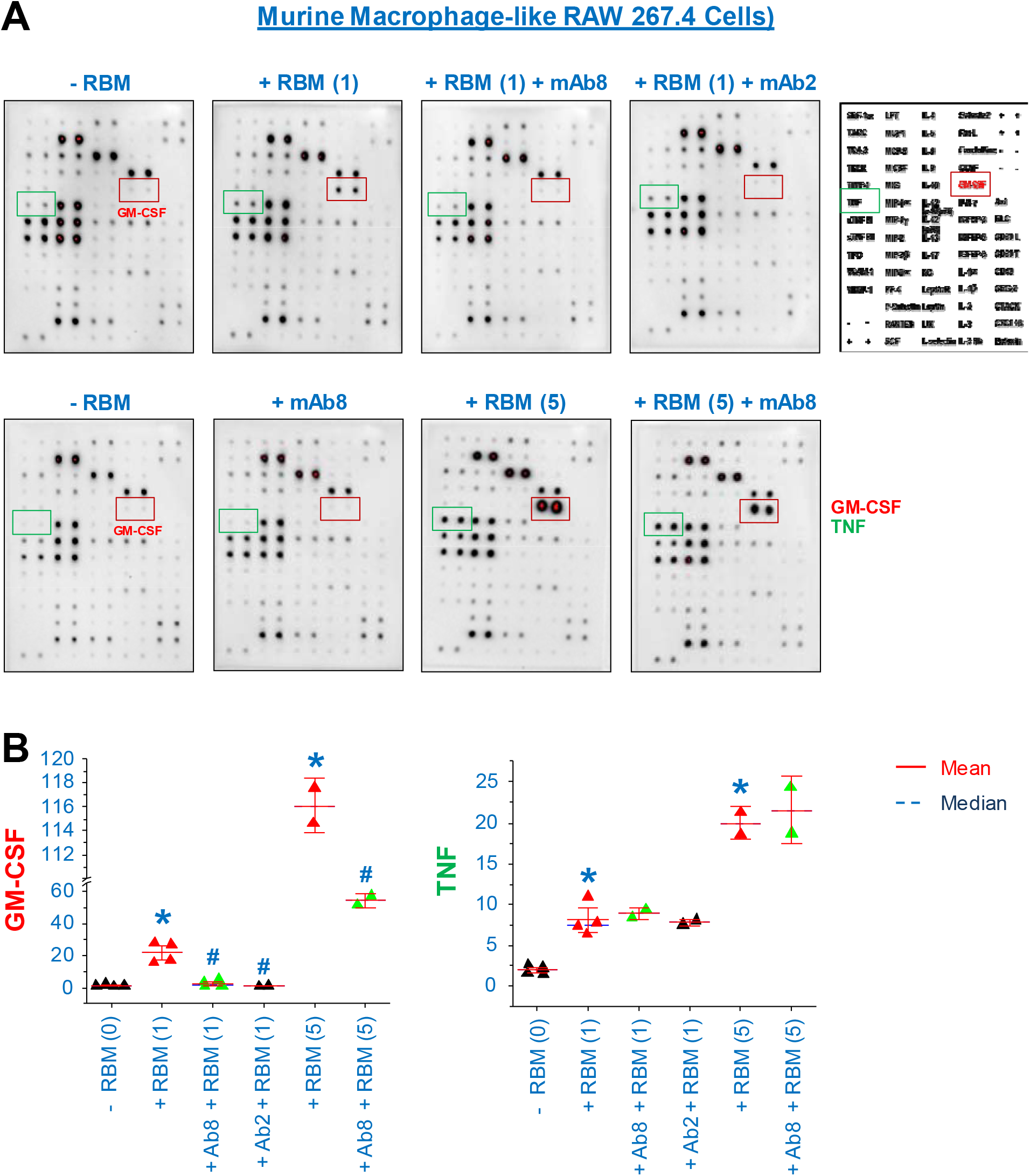
RBM-reactive mAbs blocked the RBM-induced GM-CSF secretion in murine macrophage-like RAW 264.7 cells. Murine macrophage-like RAW 264.7 cells were stimulated with recombinant RBM (1.0 or 5.0 μg/ml) either alone or in the presence of two different RBM-binding mAbs (at a molar ratio of 1:6), and extracellular concentrations of 62 cytokines and chemokines were measured by Cytokine Antibody Arrays at 16 h post stimulation. *, *P* < 0.05 versus negative controls (“-RBM” or “-R”); #, *P* < 0.05 versus positive controls (“+ RBM” or “+R”) at respective concentrations.

## Discussion

In the present study, we have generated a highly purified recombinant RBM corresponding to residue 437-508 of SARS-CoV-2, and confirmed its well-preserved ACE2-binding properties. Furthermore, we have found two RBM-cross-reactive monoclonal antibodies that competitively inhibited RBM-ACE2 interaction and selectively inhibited the RBM-induced GM-CSF secretion in both human monocyte and murine macrophage cultures. It raised an exciting possibility that vaccines capable of eliciting RBM-targeting antibodies may similarly attenuate the SARS-CoV-2-induced GM-CSF production and “cytokine storm” in clinical settings.

“Cytokine storm” refers a hyperactive inflammatory response manifested by the excessive infiltration, expansion and activation of myeloid cells (e.g., monocytes and macrophages) and consequent production of various cytokines and chemokines (e.g., GM-CSF, TNF, IL-1β, IL-6, and MCP-1). It has also been suggested as a “driver” of the disease progression particularly in a subset (~ 20%) of COVID-19 patients with more severe pneumonia that often escalates to respiratory failure and death ^26–29^. Furthermore, GM-CSF might also be a key mediator of the cytokine storm in COVID-19 and other inflammatory diseases ^30, 31^. First, GM-CSF was upregulated before TNF, IL-6, and MCP-1 in animal model of SARS-CoV infection ^32^, and its excessive production adversely contributed to the SARS-CoV-induced lung injury ^32^. Second, consistent with the critical contribution of myeloid cells to cytokine storm ^28^, the percentage of GM-CSF-expressing leukocytes was significantly increased in a subset of patients with severe COVID-19 ^33, 34^. Thus, the excessive production of GM-CSF may adversely propagate a dysregulated cytokine storm in a subset of COVID-19 patients (**Fig. 6**). One the one hand, GM-CSF can promote myelopoiesis by mobilizing progenitor myeloid cells to sites of SARS-CoV-2 infection, and facilitating their proliferation and differentiation into various innate immune cells, such as monocytes, macrophages and dendritic cells ^30^. On the other hand, GM-CSF can also polarize mature myeloid cells into a pro-inflammatory phenotype, and stimulate the production of various proinflammatory cytokines (e.g., TNF, IL-1β and IL-6) and chemokines (e.g., MCP-1) ^30^.

**Figure 6.**
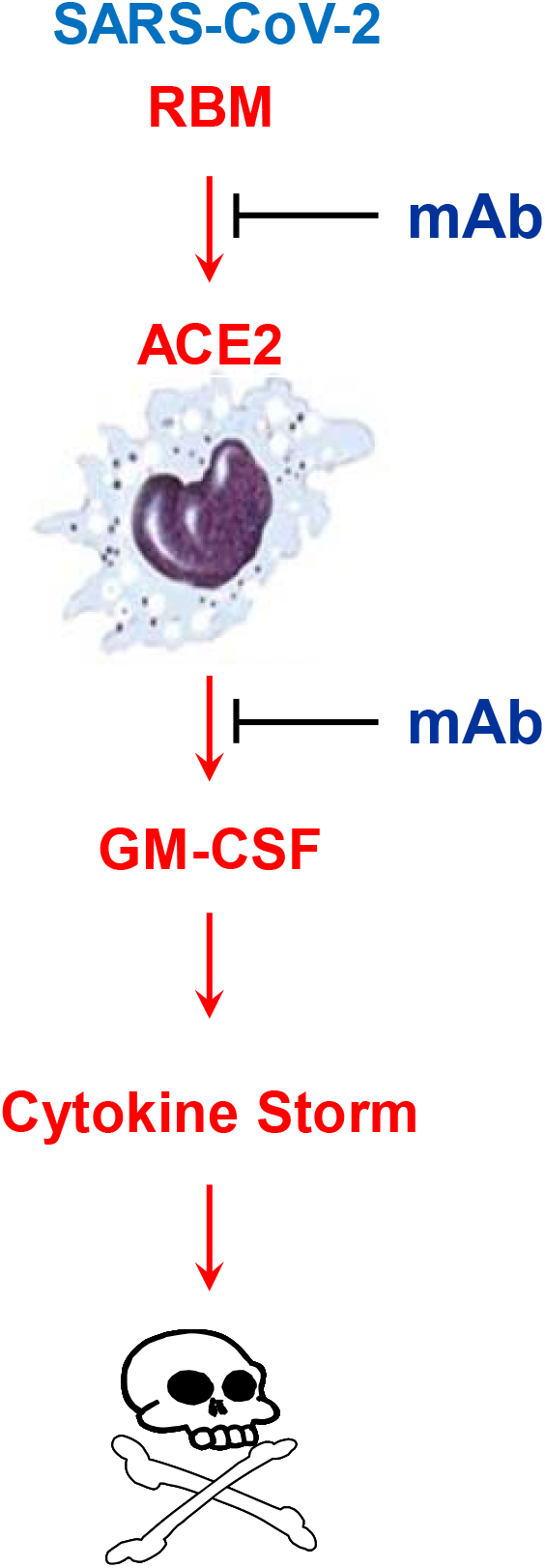
Proposed model for the mAb-mediated inhibition of SARS-CoV-2 RBM-induced GM-CSF secretion. SARS-CoV-2 RBM may bind ACE2 receptor to trigger the specific secretion of GM-CSF by macrophages and monocytes. Monoclonal antibodies capable of interrupting RBM-ACE2 interaction impairs the RBM-induced GM-CSF secretion without affecting the RBM-induced release of other pro-(e.g., IL-1β, IL-6, TNF) and anti-inflammatory cytokines (e.g., IL-10) or chemokines (MCP-1 and MIP-1δ). It might be important to assess the innate immune-modulating properties of antibodies raised against various future SAR-CoV-2 vaccines.

Currently, GM-CSF has attracted substantial interest as a therapeutic target for the clinical management of COVID-19 ^31, 35^. For instance, several companies were actively planning for COVID-19 clinical trials using anti-GM-CSF monoclonal antibodies (Clinical Trial Registry #: NCT04341116, NCT04351243, NCT04351152, NCT04376684) ^31, 35, 36^. In a recent clinical study, repetitive intravenous infusion of an anti-human GM-CSF mAb (Lenzilumab, 600 mg, thrice) significantly improved blood oxygenation, and simultaneously reduced blood levels of two pro-inflammatory cytokines (e.g., IL-1α and IL-6) in 11 out of 12 patients with severe COVID-19 ^37^. It will thus be important to verify whether our GM-CSF-inhibiting mAbs (mAb8 or mAb2) are similarly protective against SARS-CoV-2 infection in experimental and clinical settings. This is relevant because our mAbs have already been proven protective in animal model of sepsis partly by reversing immunosuppression and enhancing anti-microbial immune responses ^23^. In light of the on-going effort in developing effective SARS-CoV-2 vaccines, it may be important to assess the innate immune-modulating properties of all vaccine candidates and respective antibodies in experimental and clinical settings ^38^.

Currently, it is not known whether COVID-19 patients with more severe disease progression are generally deficient in RBM-reactive antibodies? Similarly, it is not clear whether different vaccines elicit distinct antibodies that may divergently affect SARS-CoV-2-induced cytokine storm and organ dysfunction? Finally, it remains elusive whether some SARS-CoV-2-elicited antibodies also cross-react with host proteins in such a way that may adversely compromise the half-life and/or efficacy of some protective antibodies? However, our present study has suggested a novel strategy to prevent SARS-CoV-2-elicited “cytokine storm” using RBM-targeting antibodies. Moreover, we have provided an experimental reagent (RBM) and an innate immune cell-based assay for the on-going investigation of the complex pathophysiology of COVID-19 as well as evaluation of the protective efficacy and innate immune-modulating properties of various SARS-CoV-2 vaccines.

## Materials and Methods

### Material

Murine macrophage-like RAW 264.7 cells were obtained from American Type Culture Collection (ATCC, Rockville, MD). Dulbecco’s modified Eagle medium (DMEM, 11995-065) and penicillin / streptomycin (Cat. 15140-122) were from Invitrogen/Life Technologies (Carlsbad, CA). Fetal bovine serum was from Crystalgen (FBS-500, Commack, NY) and heat-inactivated before use. The monoclonal antibodies against human tetranectin were generated in Balb/C and C57BL/6 mice at the GenScript (Piscataway, NJ, USA) as previously described ^23^. Highly purified recombinant human ACE2 corresponding to the extracellular domain (Gln18-Ser740) was obtained from two different commercial sources, Biolegend (Cat. # 7920008) and Raybiotech (Cat. # 230-30165).

### Cell culture

Human blood was purchased from the New York Blood Center (Long Island City, NY, USA), and human peripheral blood mononuclear cells (HuPBMCs) were isolated by density gradient centrifugation through Ficoll (Ficoll-Paque PLUS) as previously described ^23^. Murine macrophages or human monocytes (HuBPMCs) were cultured in DMEM supplemented with 1% penicillin/streptomycin and 10% FBS or 10% human serum. When they reached 70-80% confluence, adherent cells were gently washed with, and immediately cultured in, OPTI-MEM I before stimulating with highly purified recombinant RBD or RBM in the absence or presence of anti-TN mAbs. The extracellular concentrations of various cytokines/chemokines were determined by Cytokine Antibody Arrays as previously described ^39^.

### Preparation of recombinant RBD and RBM proteins

The cDNAs encoding for the ACE2 receptor binding domain (RBD, residue 319 - 541) or receptor binding motif (RBM, residue 437-508) of SARS-CoV-2 spike protein (S) were cloned into a pCAL-n vector, and the recombinant proteins with an N-terminal Histidine Tag (6 × His) were expressed in *E. coli* BL21 (DE3) cells in the presence of 3.0 mM IPTG (isopropyl-1-thio-beta-D-galactopyranoside). Recombinant RBD and RBM proteins were isolated from the inclusion bodies by differential centrifugation, and further purified by urea (8.0 M Urea, 20 mM Tris-HCl, pH 8.9) solubilization and agarose bead-immobilized metal (Ni^2+^) affinity chromatography. After extensive washing with buffer 1 (20 mM Tris-HCl, 10 mM imidazole, 0.5 M NaCl, 8.0 M Urea, pH 8.0) and buffer 2 (20% DPBS1X, 10% glycerol, 8.0 M Urea, pH 7.5), the recombinant histidine-tagged RBD or RBM proteins were eluted with buffer containing 0.5 M Imidazole,10% Glycerol, 20% DPBS1X, 8.0 M Urea, pH 8.0. The recombinant proteins were then further purified by dialysis at 4⁰ C in buffer containing 20% DPBS1X, 10 % Glycerol and 0.5 mM TCEP, pH8.0. Recombinant proteins were tested for LPS content by the chromogenic *Limulus* amebocyte lysate assay (Endochrome; Charles River), and the endotoxin content was less than 0.01 U per microgram of recombinant proteins.

### Open Surface Plasmon Resonance (SPR)

We used the Nicoya Lifesciences gold-nanoparticle-based Open Surface Plasmon Resonance (OpenSPR) technology to estimate the binding kinetics and affinity of ACE2 or monoclonal antibodies to SARS-CoV-2 RBD or RBM following the manufacturer’s instructions. For instance, highly purified recombinant RBD or RBM was immobilized on the NTA sensor chip (Cat. # SEN-Au-100-10-NTA), and ACE2 or mAb was applied at different concentrations. The response units were recorded over time, and the binding affinity was estimated as the equilibrium dissociation constant K_D_ using the Trace Drawer Kinetic Data Analysis v.1.6.1. (Nicoya Lifesciences) as previously described ^23^. To determine the possible competition with the human ACE2, SARS-CoV-2 RBM was immobilized to NTA sensor chips via histidine tag for a final RU around 500. A RBM-binding mAb was injected onto the chip until binding steady-state was reached, and ACE2 was re-injected as analyte at identical concentrations. The competition capacity of RBM-binding mAb was determined by the level of reduction in response units of ACE2 with and without prior mAb incubation. Results presented are representatives of two independent experiments.

### Cytokine Antibody Array

Human Cytokine Antibody C3 Arrays (Cat. No. AAH-CYT-3-4), which detect 42 cytokines on one membrane, were used to determine cytokine concentrations in human monocyte-conditioned culture medium as previously described ^23^. Murine Cytokine Antibody Arrays (Cat. No. M0308003, RayBiotech Inc.), which simultaneously detect 62 cytokines on one membrane, were used to measure relative cytokine concentrations in macrophage-conditioned culture medium as described previously ^23, 40^.

### Statistical analysis

All data were assessed for normality by the Shapiro-Wilk test before conducting statistical tests among multiple groups by one-way analyses of variance (ANOVA) followed by the Fisher Least Significant Difference (LSD) test. A *P* value < 0.05 was considered statistically significant.

## ACKNOWLEDGEMENTS

This work was supported in part by the National Institutes of Health (NIH) grants R01GM063075, and R01AT005076.

## AUTHOR CONTRIBUTIONS

X.Q. performed innate immune cell-based experiments; S.Z. performed the Open SPR experiments; J.L. and K.J.T. generated the recombinant RBM and RBD proteins; P. W. provided important input to the experimental design. H.W. supervised the study, interpreted the results, and wrote the manuscript.

